# Loss of ASIC1A-dependent inhibitory neuron activity in basolateral amygdala is associated with increased CO_2_-evoked jumping

**DOI:** 10.64898/2026.05.18.725939

**Authors:** Rebecca J Taugher-Hebl, Aubrey C Chan, Collin J Kreple, Ali Ghobbeh, Grace Z Wang, Gail I S Harmata, Mackenzie M Conlon, Subhash C Gupta, Rong Fan, Ramkumar Kuruba, Margaret P Price, Jeffrey D Long, Youngcho Kim, Brian J Dlouhy, Nandakumar S Narayanan, John A Wemmie

## Abstract

**Background:** Responding appropriately to threats is critical for survival. Dysregulated defensive responses are core features of psychiatric illnesses including panic disorder and post-traumatic stress disorder. Carbon dioxide (CO_2_) inhalation evokes defensive behaviors in both humans and mice. Here we investigated the role of acid-sensing ion channels (ASICs) in CO_2_-evoked jumping in mice.

**Methods:** Defensive behaviors (jumping and freezing) were assessed in response to CO_2_ inhalation and basolateral amygdala (BLA) acidification. We tested the role of ASICs using global knockout mice and *Asic1a^loxP/loxP^* mice transduced with AAV-CMV-Cre or AAV-CaMKII-Cre in the BLA. Effects of CO_2_ on single neuron firing and local field potentials were studied via BLA microwire arrays.

**Results:** ASIC1A disruption increased CO_2_-evoked jumping while reducing freezing, paralleled by increased BLA c-Fos induction. Acidification of the BLA recapitulated these effects. Virus-mediated ASIC1A disruption in BLA did not resolve the locus of ASIC1A action in jumping. CO_2_ inhalation suppressed firing in most BLA neurons, though a small number increased firing. ASIC1A disruption enhanced CO_2_-induced suppression of narrow waveform neurons (putative interneurons), and facilitated excitation of wide waveform neurons (putative principal neurons). Additionally, CO_2_ produced concentration-dependent broadband power suppression with selective theta enhancement, effects that were augmented by ASIC1A disruption.

**Conclusions:** Together, these findings suggest that ASIC1A promotes interneuron activity during acidosis and that its loss may reduce inhibition of principal neuron output, shifting defensive responses from freezing toward jumping. These results advance our understanding of how brain pH and ASICs regulate defensive behavior, with potential implications for understanding dysregulated defensive responses.

## Introduction

Responding appropriately to threats is critical for survival. Humans and other animals need to quickly evaluate the proximity and intensity of danger and initiate suitable defensive strategies. Responses to threats can be diverse and exist on a spectrum whereby the best defensive actions change as the level of risk increases or decreases [1, 2]. Arousal or avoidance responses may be best when danger is relatively low, whereas fleeing or fighting responses may be best when danger is high and the threat is imminent. Defensive responses that are out of proportion to the level of threat can also be highly maladaptive and disruptive, for example in psychiatric illnesses such as post-traumatic stress disorder (PTSD) and panic disorder.

A useful tool for studying defensive responses is carbon dioxide (CO_2_). CO_2_ is produced by cellular respiration and is normally maintained within a narrow physiological range, largely through breathing [3]. Rising CO_2_ levels herald the rapidly lethal threat of suffocation: the higher the CO_2_ concentration, the more immediate the threat. In a laboratory setting, CO_2_ inhalation can be used to evoke a range of defensive responses depending on the dose and duration of exposure. In humans, inhaling CO_2_ can elicit increased anxiety, fear, and panic – especially in individuals with panic disorder [4–8]. Similarly, in mice, CO_2_ evokes a range of defensive behaviors including avoidance, freezing, and jumping [9–12].

We have identified a key role for the amygdala in CO_2_ responses in both humans and mice. When given a CO_2_ inhalation challenge, humans with bilateral lesions of amygdala (including basolateral amygdala (BLA)) exhibited increased panic, but did not develop anticipatory anxiety in response to subsequent CO_2_ challenges [13], suggesting the amygdala is required for learned anticipatory responses but suppresses acute panic responses to CO_2_. In mice, bilateral lesions of the BLA reduced CO_2_-evoked freezing and increased CO_2_-evoked jumping [11]. Taken together, these observations suggest that the amygdala plays a critical role in regulating CO_2_-evoked defensive responses.

CO_2_ inhalation induces acidosis, which is likely responsible for molecular and cellular effects contributing to defensive behaviors. Our work has found an important role for acid-sensing ion channels (ASICs) in defensive responses to CO_2_ [9, 10, 14]. ASICs are cation channels activated by extracellular acidosis. These channels are expressed on the neuronal plasma membrane at both the soma and in dendritic spines, and have been implicated in pathological effects of acidosis [15, 16] as well as synaptic plasticity and neurotransmission [17–19]. These channels are formed by trimeric assemblies of subunits, with ASIC1A and ASIC2A subunits being important for determining pH sensitivity, desensitization kinetics, and Ca^2+^ permeability [17, 20]. Both ASIC1A and ASIC2A contribute to synaptic ASIC currents [19], are expressed widely in the brain, and are relatively abundant in regions known to be critical for defensive responses such as the BLA [14, 21]. Previously, we found that disrupting ASIC1A or ASIC2 in mice reduced avoidance and freezing responses to CO_2_ [9, 10, 14], although jumping was not assessed at the time. Given the robust expression of ASIC1A and ASIC2 in the BLA and our subsequent work implicating the amygdala in CO_2_-evoked jumping [11], here we tested the hypothesis that ASICs in the BLA also contribute to this high-intensity defensive response.

## Methods

### Mice

All mice were maintained on a congenic C57BL/6 background. *Asic1a^-/-^*[22], *Asic2^-/-^* [23], *Asic1a^loxP/loxP^* [10, 19], and *Syn Asic1a* KO mice [24] were generated and maintained as previously described. Experimental groups were sex- and age-matched, with mice 12-20 weeks old being used for behavioral studies. All experiments were conducted during the light cycle. The University of Iowa Animal Care and Use Committee approved all experiments.

### Gas Exposure for Behavioral Analysis

Mice were exposed to different concentrations of CO_2_ as previously described [9–11, 24]. In brief, compressed air, 10% CO_2_ (10% CO_2_, 21% O_2_, balanced with N_2_), and 20% CO_2_ (20% CO_2_, 21% O_2_, balanced with N_2_) gas mixtures were infused into a custom-made Plexiglas chamber at a rate of 5L/minute to pre-fill the chamber with the specified gas mixture. Mice were placed into the chamber for 10 minutes, during which behavior was videotaped. A trained observer blinded to condition and genotype assessed jumping and freezing. Jumping was defined as all four paws simultaneously being airborne and freezing was defined as an absence of motion other than breathing.

### c-Fos Immunohistochemistry

Mice were exposed to compressed air or 10% CO_2_, as described above. 1 hr and 15 minutes after the start of the gas exposure, mice were perfused with 4% PFA and brain tissue was harvested for analysis and 40 μm sections were taken through the BLA. c-Fos immunohistochemistry was performed using the Fast Fos method [25]. Reagents used were: Streptavidin/Biotin Blocking Kit (Vector Biolabs, SP-2002), horse serum (Jackson ImmunoResearch Laboratories Inc., 008-000-121), c-Fos goat polyclonal antibody (Santa Cruz, SC52G), biotinylated horse anti-goat IgG (Vector Biolabs, BA-9500), and streptavidin-HRP complex (Jackson ImmunoResearch Laboratories Inc., 016-030-084). DAB staining was performed using a DAB Peroxidase Substrate Kit (Vector Biolabs, SK-4100). Brain slices were mounted on slides and imaged using an Olympus BX-61 microscope.

### Low pH injections

Guide cannulae were implanted in the amygdala in a manner similar to that which has been previously described [9, 10, 26]. In brief, guide cannulae were implanted in the amygdala and a 23-gauge guide cannula (Plastics One) was implanted dorsal to the amygdala (1.4 mm posterior from bregma, 3.2 mm lateral to the midline, and 2.9 mm ventral to the pial surface). After a minimum of three days recovery, 1 μl of acidic (pH 3.0) or normal pH (7.3) ACSF was infused into the amygdala by inserting a 30-gauge injector (Plastics One) 1 mm beyond the tip of the guide cannula into the amygdala. We have previously validated this technique and found that it induces a focal acidosis of pH ∼6.8, there is no tissue injury, and no acidosis is detected in other nearby brain sites [9, 10]. Immediately following the injection, mice were placed in a custom-made Plexiglas chamber for 10 minutes, during which they were videotaped. A trained observer blinded to condition and genotype assessed freezing and jumping in the same manner as during gas exposure. Targeting of cannulae was confirmed postmortem via methylene blue infusion.

### Virus injections

AAV2/1-CMV-eGFP and AAV2/1-CMV-Cre were purchased from the University of Iowa Gene Transfer Vector Core. AAV1-CaMKII-Cre (cat# 105558-AAV1) and AAV1-CaMKII-eGFP (cat# 105541-AAV1) were purchased from Addgene. AAVs were stereotactically injected into BLA (1.4 mm posterior from bregma, 3.2 mm lateral to the midline, and 3.9 mm ventral to the pial surface) or BNST (0.4 mm anterior to bregma, 1.0 mm lateral to the midline, and 4.3 mm ventral to the pial surface) of *Asic1a^loxP/loxP^*mice bilaterally, in a manner similar to that previously described [10, 19, 27]. 0.5 μl of virus was infused into each site. Behavioral testing was performed at least 3 weeks after surgery to allow time for recovery and viral transduction. Targeting of injections was verified postmortem and bilateral transduction in the structure of interest (BLA or BNST) was required to be considered a hit.

### Western blotting

Western blotting of BLA protein lysates was performed as previously described [10, 24, 28]. In brief, lysates were obtained from 1.2 mm punches of the BLA taken from 2 mm coronal sections. Primary antibodies used were rabbit polyclonal anti-ASIC1 antiserum [21] diluted 1:1,000 and chicken polyclonal anti-GAPDH, diluted 1:10,000. Secondary antibodies used were donkey anti-rabbit IRDye 800 CW, diluted 1:10,000 and donkey anti-chicken IRDye 680 LT, diluted 1:20,000 (LI-COR). Membranes were imaged with an Odyssey imaging system (LI-COR) and intensity of bands was measured using ImageJ.

### Acid-evoked currents

*Asic1a^loxP/loxP^* mice were injected with AAV2/1-CMV-eGFP or a mixture of AAV2/1-CMV-eGFP and AAV2/1-CMV-Cre, as described above. After allowing at least 3 weeks for viral transduction, coronal slices were made through the BLA. Slices were cut, and whole-cell recordings were performed as previously described [28]. Transduced neurons were identified by their eGFP expression. Putative BLA principal neurons were identified based on their morphology and membrane capacitance. The patch solution was potassium gluconate-based and a −70 mV holding potential was used. Recordings were performed in presence of CNQX, picrotoxin, and AP5. Acid-evoked currents were elicited by applying pH 5.6 for 3 seconds. pH 5.6 was chosen as a maximal stimulus to confirm the presence or absence of functional ASIC1A channels, rather than to approximate physiological acidosis. Current density (peak amplitude / membrane capacitance) is reported.

### Open Field

Locomotor activity was assessed in an open field chamber (40.6 cm wide x 40.6 cm deep x 36.8 cm high, San Diego Instruments) for 30 min as previously described [10].

### In vivo neural recordings

Mice were stereotactically implanted with custom 16-channel microwire arrays (Microprobes, Gaithersburg, MD) targeting the BLA (1.4 mm posterior from bregma, 3.4 mm lateral to the midline, and 3.9 mm ventral to the pial surface). Each mouse was implanted unilaterally, counterbalanced across the cohort so both left and right BLA were represented. Microwire arrays were affixed to the skull with bone anchor screws, cyanoacrylate glue, and dental cement. Mice were allowed to recover from surgery for at least 9 days and acclimated to handling for at least 5 minutes each day on post-op days 5-9. Recordings took place in a custom-built acrylic chamber; the exposure paradigm consisted of a 10-minute baseline period followed by three epochs for compressed air, 10% CO_2_, and 20% CO_2_. Each epoch consisted of 15 minutes of the gas then 15 minutes in room air before the next epoch. Gas exposures were performed in a single session to minimize disturbances to the head cap from connection and disconnection. Continuous recordings were obtained at 40 kHz and field potential recordings were obtained at 1000 Hz using Plexon equipment and software. Location of the microwire arrays was verified postmortem. 14 μm sections were taken through the amygdala and Nissl stained. Slides were imaged with an Olympus BX61VS microscope and animals in which the MWA did not hit the amygdala were excluded from the analysis.

### Single neuron data analysis

Single neuron spikes were analyzed as previously described [29]. Briefly, continuous data was imported into MATLAB, filtered, then referenced using truncated means from combined channels. Spikes were included only if they exceeded an amplitude threshold of 4.5x median absolute deviation. Individual waveforms were then sorted manually in Offline Spike Sorter software (Plexon). Waveforms were sorted based on clustering in principal component analysis space, waveform shape, and interspike interval distribution. Sorted waveforms were imported into MATLAB software for further analysis of firing frequency and action potential width. Action potential width was based on trough-to-peak time, with 550 μs as the cutoff between narrow and wide waveforms [30]. Firing frequency was calculated using 5-second bins and z-normalized to mean firing frequency during the room air baseline period. A neuron was considered to increase firing during an epoch if its average z-score over that epoch exceeded 1, and to decrease firing during an epoch if its average z-score was below -1.

### Field potential data analysis

Field potential recordings were extracted from raw data files using NeuroExplorer software (Plexon) and analyzed using custom code in MATLAB. Raw LFP signal was averaged across 16 channels for each animal, then power at different frequencies was obtained by wavelet convolution across 75 frequency steps (1-20 Hz in 1 Hz steps, then 22-130 Hz in 2 Hz steps) [31]. Power data was normalized to the average power between -8 min to -2 min of each gas’s pre-exposure period, then averaged into 1 second bins. Power in each frequency band was obtained by taking the mean of all frequencies belonging in that band (delta: 1-4 Hz, theta: 5-8 Hz, alpha: 9-12 Hz, beta: 13-30 Hz, gamma: 32-48 Hz, high gamma: 72-110 Hz). Frequencies ranging from 50-70 were not analyzed to eliminate possible effects of 60 Hz line noise. For purposes of plotting, power was further averaged into 30 second bins.

### Statistical analysis

Statistical analyses were performed in Prism (Graphpad), MATLAB (MathWorks), SPSS (IBM), and Excel (Microsoft). p < 0.05 was considered significant for all analyses. Data are plotted as mean ± SEM. Differences between two groups were tested with a Student’s t-test if the variable was continuous (e.g. time freezing) or with a Mann-Whitney test if the variable was discrete (e.g. number of jumps). In the context of a Student’s t-test, an F-test was performed to determine if variances were significantly different, and a Welch’s correction was used when appropriate. Differences between more than two groups were assessed with an ANOVA with planned contrast testing in instances in which differences in groups were hypothesized *a priori*. For the field potential analysis, power at each frequency band was analyzed with mixed effects models incorporating time, genotype, gas exposure, and their interactions as fixed effects, and mouse as a random effect using MATLAB. For individual spike firing, distributions of neuron responses were compared using chi-squared tests where possible, or Fisher’s exact test if the distributions did not meet criteria for using a chi-squared test (calculated expected values less than 1, or less than 20% of expected values were greater than 5). Comparison of changes in firing frequency was performed using the Mann-Whitney test. Comparison of changes in firing frequency across genotypes and gas conditions was performed using linear mixed models analysis.

## Results

To understand the role of ASICs in jumping responses to CO_2_, we exposed wild-type, *Asic1a^-/-^*, and *Asic2^-/-^* mice to a range of CO_2_ concentrations (compressed air, 10% and 20% CO_2_) capable of eliciting freezing and jumping in wild-type mice [11]. Both *Asic1a^-/-^* and *Asic2^-/-^*mice displayed robust jumping in 10% CO_2_ that was rare in wild-type mice (**Figure 1A**). When CO_2_ concentration was increased to 20%, wild-type mice began to display robust jumping. *Asic2^-/-^*mice displayed similar levels of jumping in 20% CO_2_ as compared to wildtypes, while *Asic1a^-/-^*mice displayed a further marked increase in jumping at 20% CO_2_. Consistent with our previous observations [9, 10, 14, 24], freezing was robustly evoked by 10% CO_2_ in wild-type mice and was markedly decreased in *Asic1a^-/-^* and *Asic2^-/-^*mice (**Figure 1B**). Freezing was increased in 20% CO_2_ as compared to 10% CO_2_ in mice of all three genotypes, with *Asic1a^-/-^* and *Asic2^-/-^*mice still exhibiting less freezing than wild-type controls. As expected, wild-type mice exhibited minimal jumping and freezing in compressed air (**Figures 1A**, **1B**). Both *Asic1a^-/-^* and *Asic2^-/-^*mice displayed minimal freezing in compressed air and low levels of jumping, though *Asic1a^-/-^*mice did jump more than wild-type controls (**Figure 1A**). Together, these observations suggest that ASICs oppose CO_2_-evoked jumping behavior and promote CO_2_-evoked freezing. The effects of ASIC1A and ASIC2 disruption on freezing were largely similar. However, the more pronounced jumping phenotype with ASIC1A disruption suggests that the mechanisms underlying freezing and jumping are at least partially separable. Given our interest in jumping responses, we subsequently focused on better understanding the role of ASIC1A.

**Figure 1.**
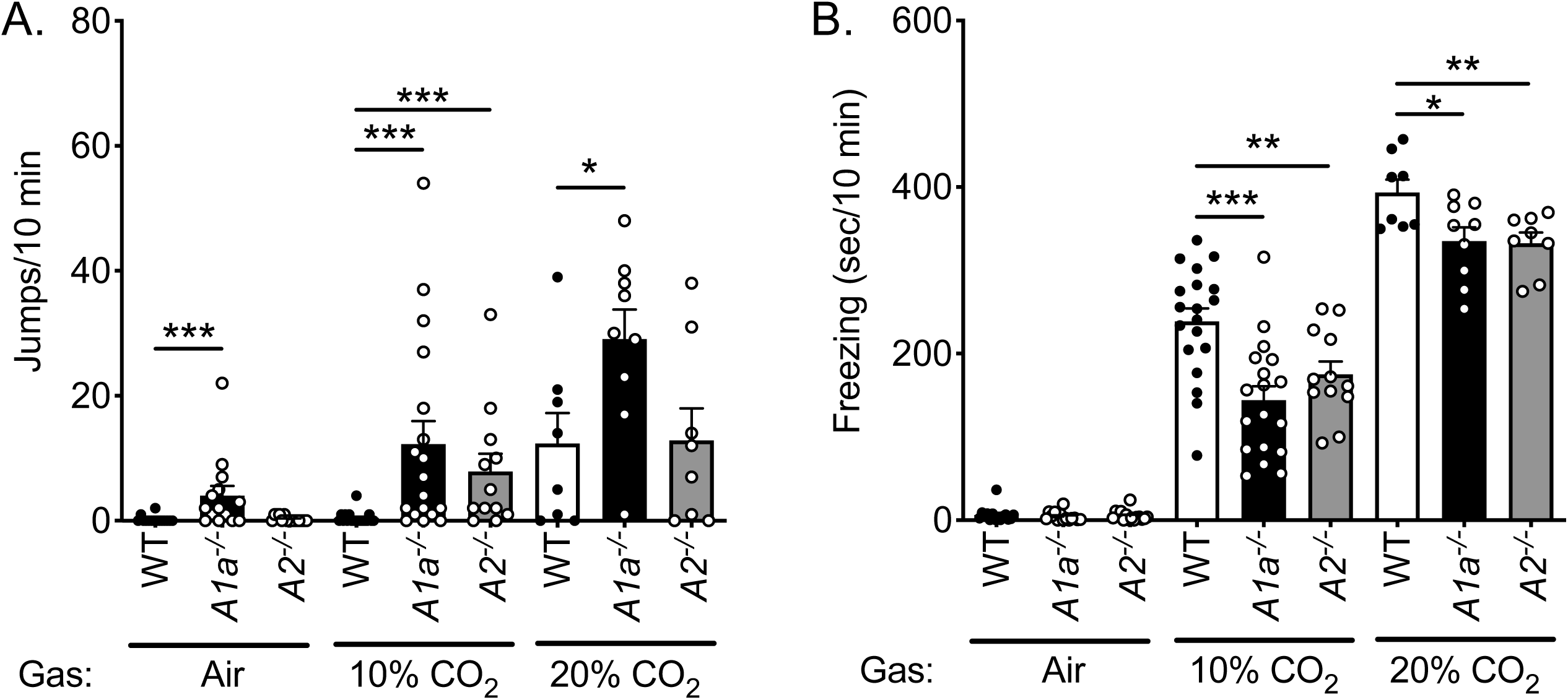
CO_2_-evoked jumping depends on acid-sensing ion channels. **A)** A comparison of jumping responses in wild-type (WT), *Asic1a^-/-^* (*A1a^-/-^*) and *Asic2^-/-^* (*A2^-/-^*) mice exposed to increasing concentrations of CO_2_ (Air, 10%, and 20% CO_2_) revealed effects of both genotype (p < 0.0001, n = 8-19 per group) and CO_2_ concentration (p < 0.0001) as well as a trend towards a genotype by CO_2_ concentration interaction (p = 0.0703). *Asic1a^-/-^* mice jumped more than wild-type controls in compressed air (*** p = 0.0008), 10%CO_2_ (*** p < 0.0001), and 20% CO_2_ (*p = 0.0294), whereas *Asic2^-/-^* displayed similar levels of jumping as wild-type controls in compressed air (p = 0.6513) and 20% CO_2_ (p = 0.9386), but elevated jumping to 10% CO_2_ (*** p < 0.0001). **B)** Quantification of freezing responses across CO_2_ concentrations revealed a genotype by CO_2_ concentration interaction (p = 0.0023). Little freezing was observed in compressed air, whereas *Asic1a^-/-^* and *Asic2^-/-^* mice froze less than wild-type controls in both 10% (*Asic1a^-/-^*: ***p = 0.0002, *Asic2^/-^*: *p = 0.0093) and 20% CO_2_ (*Asic1a^-/-^*: *p = 0.0216, *Asic2^-/-^*: **p = 0.0093).

To test our hypothesis that ASICs in the BLA oppose CO_2_-evoked jumping, we quantified induction of c-Fos protein expression in BLA after exposure to 10% CO_2_ vs. compressed air. CO_2_ increased c-Fos induction in BLA in both wild-type and *Asic1a^-/-^* mice (**Figure 2A**, **2B**). *Asic1a^-/-^*mice exhibited greater c-Fos induction than wild types in both compressed air and 10% CO_2_, with a significant gas × genotype interaction reflecting a particularly large increase in *Asic1a^-/-^* mice exposed to CO_2_ (**Figure 2B**). Though our c-Fos labeling did not distinguish between specific neuron populations, it is likely that the increased c-Fos signal reflects, at least in part, increased principal neuron activation, given that these cells constitute the large majority of BLA neurons and carry output to downstream sites. Therefore, the increased c-Fos induction in *Asic1a^-/-^* mice parallels the increase in jumping and suggests that loss of ASIC1A leads to a relative increase in BLA activity which may contribute to the shift in defensive behavior towards jumping.

**Figure 2.**
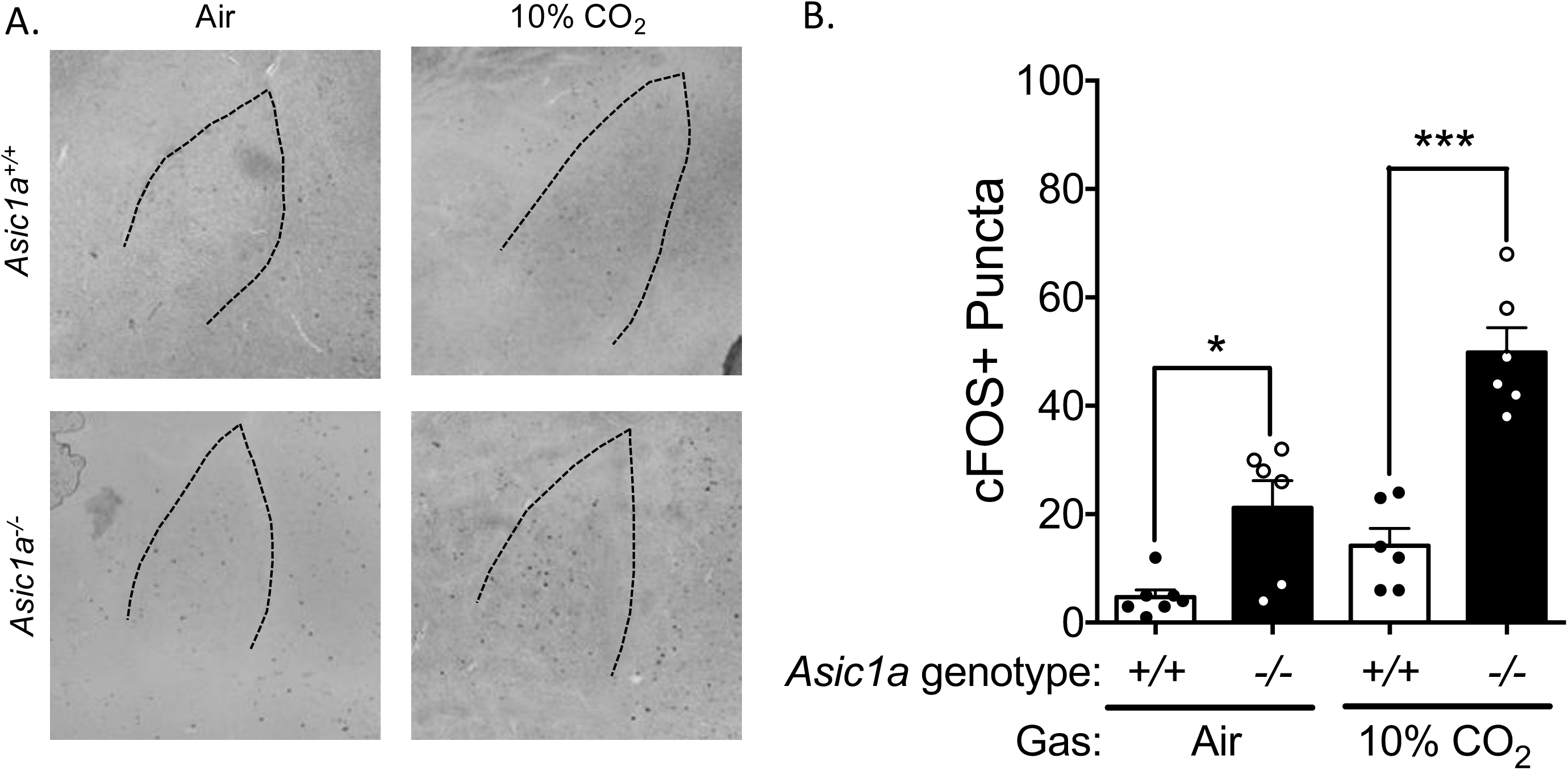
10% CO_2_ inhalation induces ASIC1A-dependent c-Fos expression in the BLA. **A)** Representative c-Fos expression in the BLA 1 hr and 15 min after exposure to compressed air or 10% CO_2_ in *Asic1a^+/+^* and *Asic1a^-/-^* mice. **B)** CO_2_ induced c-Fos expression in an ASIC1A-dependent manner (genotype x treatment interaction: F(1, 21) = 6.776, p = 0.0166, n = 6-7). Relative to *Asic1a^+/+^* controls, *Asic1a^-/-^* mice exhibited increased BLA c-Fos both at baseline (*p = 0.0210) and after exposure to 10% CO_2_ (***p < 0.0001).

To probe whether jumping was due to CO_2_-induced acidosis or some other effect of CO_2_, we injected neutral or acidic artificial cerebrospinal fluid (ACSF) into the amygdala of wild-type and *Asic1a^-/-^*mice and assessed jumping and freezing behaviors (**Figure 3A**). Previously, we validated that this technique lowers brain pH to ∼6.8, similar to the acidosis seen with 10% CO_2_ inhalation and found that this infusion of acidic ACSF does not permanently damage the brain tissue and that acidosis is observed only focally at the injection site without spread to nearby brain regions [9, 10]. In this experiment, we found that acidifying the amygdala induced robust jumping in *Asic1a^-/-^*mice, but not wild-type mice (**Figure 3B**). Consistent with our previous observations [9], we found that infusion of low pH into the amygdala evoked freezing in wild-type mice that was absent in *Asic1a^-/-^* mice (**Figure 3C**). Injection of neutral pH ACSF did not evoke freezing or jumping in either wild-type or *Asic1a^-/-^* mice (**Figures 3B**, **3C**). Taken together, these observations are consistent with ASIC1A in the amygdala promoting acid-evoked freezing and opposing acid-evoked jumping, and suggest that amygdala acidosis is a critical driver of these behavioral effects during CO_2_ inhalation.

**Figure 3.**
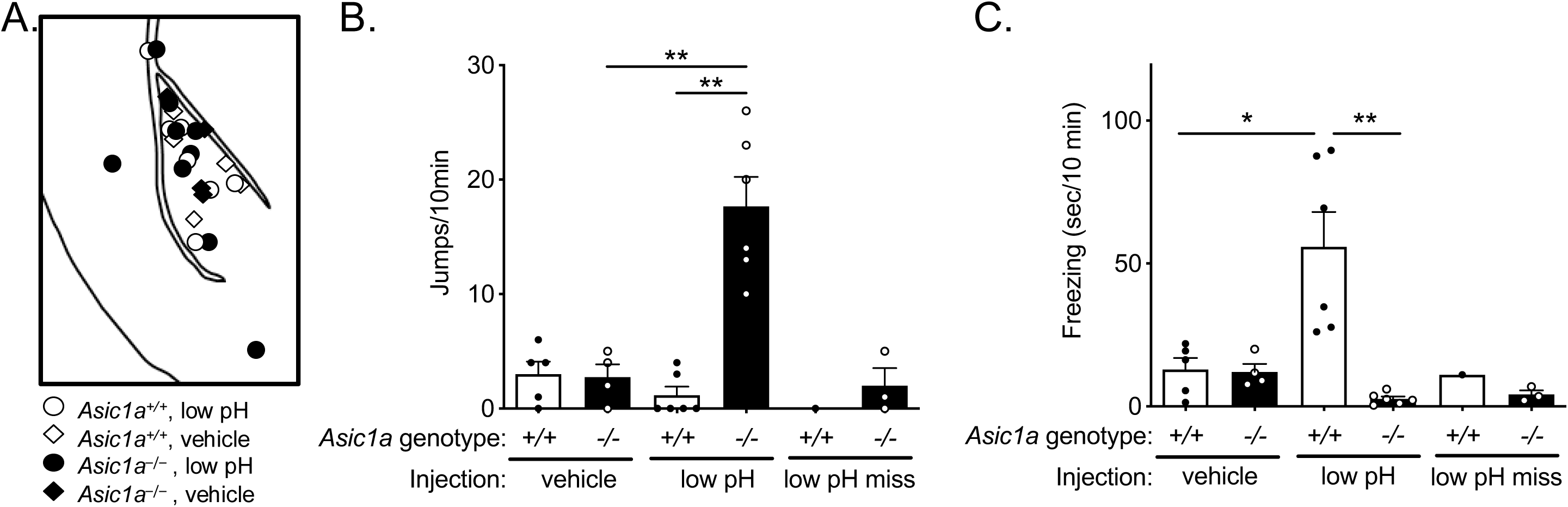
Acidification of the amygdala elicits ASIC1A-dependent freezing and jumping. **A)**Diagram of infusion site locations: low pH (circles) vs. vehicle (diamonds), wild-type (open) vs. *Asic1a^-/-^* mice (filled) **B)** Jumping behavior after infusion of low pH or normal pH solution into the amygdala. Infusion of low pH into the amygdala induced ASIC1A-dependent jumping behavior (genotype x treatment interaction: F(1, 17 = 24.07, p = 0.0001). Infusion of low pH into the amygdala evoked repeated jumping in *Asic1a^-/-^* mice that was largely absent wild-type mice (**p = 0.0022) and *Asic1a^-/-^* mice injected with a normal pH vehicle (**p = 0.0095). **C)** Freezing behavior after infusion of low pH or normal pH vehicle into the amygdala. Infusion of low pH into the amygdala induced freezing in wild-type mice that was dependent on *Asic1a* genotype (genotype x treatment interaction: F(1, 17 = 12.22, p = 0.0028, n = 4-6). Wild-type mice injected with low pH solution displayed more freezing than *Asic1a^-/-^*mice injected with low pH (**p = 0.0071) or wild-type mice injected with a normal pH vehicle solution (*p = 0.0152). Four low pH injections missed the amygdala (1 wild-type, 3 *Asic1a^-/-^*); little freezing was observed in these mice.

Because ASIC1A is widely expressed in the brain, we next sought to determine whether ASIC1A’s role in regulating acid-evoked defensive responses generalized to other brain sites. Previously, we identified the BNST as a site of ASIC1A action in CO_2_-evoked freezing and found that acidic CSF injected into the BNST was sufficient to elicit ASIC1A-dependent freezing [10], so we asked if BNST acidosis also elicits ASIC1A-dependent jumping. Though some sporadic jumping was observed, BNST acidosis did not elicit ASIC1A-dependent jumping (**Figure S1A**). In contrast, similar levels of acidosis-induced freezing were observed in wild-type mice with BNST acidosis (**Figure S1B**) and BLA acidosis (**Figure 3C**). These results are consistent with both BLA and BNST regulating freezing but suggest a more pronounced role for the BLA in jumping behavior.

We hypothesized that deleting ASIC1A specifically in the BLA would recapitulate the jumping phenotype seen in *Asic1a^-/-^* mice. To test this hypothesis, we disrupted ASIC1A in the BLA by injecting an AAV expressing Cre recombinase under control of a CMV promoter (AAV-CMV-Cre) into the amygdala of *Asic1a^loxP/loxP^* mice. We verified that this manipulation reduced ASIC1A expression in the amygdala (**Figure 4A**, **4B**) and abolished acid-evoked currents in virus-transduced BLA neurons (**Figure 4C**, **4D**), consistent with viral transduction and ASIC1A disruption of a subset of BLA neurons. Next, we assessed the effects of ASIC1A disruption on CO_2_-evoked defensive behaviors. When exposed to 10% CO_2_, neither AAV-CMV-GFP nor AAV-CMV-Cre transduced mice exhibited significant jumping (**Figure 4E**). In contrast, CO_2_-evoked freezing was attenuated in AAV-CMV-Cre transduced mice relative to AAV-CMV-GFP transduced controls (**Figure 4E)**. These observations were not explained by gross differences in motor behavior, as AAV-CMV-Cre injection did not alter locomotion in the open field (**Figure S2A**). The observation that AAV-CMV-Cre animals did not recapitulate the increase in jumping observed in *Asic1a^-/-^* animals leaves open the possibility that our viral disruption may have left residual ASIC1A activity, perhaps in a critical cell population, sufficient to suppress jumping behavior.

**Figure 4.**
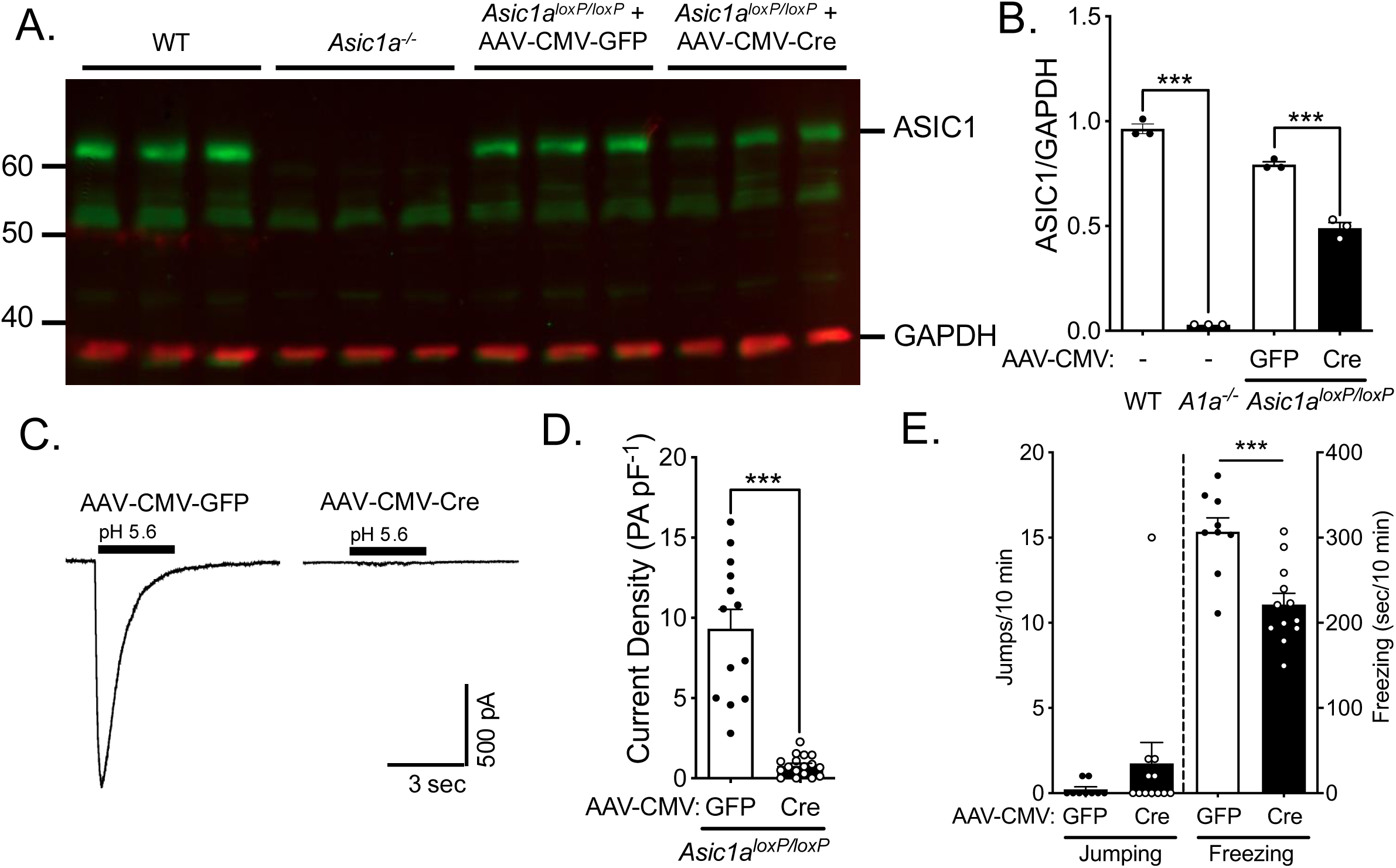
CO_2_-evoked freezing depends on ASIC1A in the amygdala. ASIC1A was disrupted in the amygdala of *Asic1a^loxP/loxP^* mice using an AAV expressing Cre recombinase (AAV-CMV-Cre). An eGFP vector (AAV-CMV-GFP) was used as a control. **A, B)** ASIC1 protein (green) is robustly expressed in amygdala tissue from wild-type mice and is absent from *Asic1a^-/-^* amygdala (t(2) = 40.00, ***p = 0.0006, n = 3/group). Injection of AAV-CMV-Cre into the amygdala reduced ASIC1 expression relative to AAV-CMV eGFP injected controls (t(4) = 1024, ***p = 0.0005, n = 3/group). GAPDH (red) was used as a loading control. **C, D)** Acid-evoked currents were largely abolished in AAV-Cre transduced neurons, but were intact in non-transduced neurons and neurons transduced with only AAV-GFP (t(12.41) = 7.093, ***p < 0.0001, n = 13, 17). **E)** Amygdala-specific disruption of ASIC1A reduced freezing to 10% CO_2_ (t(19) = 4.124, ***p = 0.0006, n = 9, 12), but did not significantly alter jumping behavior (p = 0.2561).

Because we had previously probed the role of ASIC1A in the BNST with the same AAV-CMV-Cre vector [10] and found effects on CO_2_-evoked freezing, we reanalyzed videos from those experiments to determine if jumping was also affected. We observed no CO_2_-evoked jumping following BNST-specific ASIC1A disruption (**Figure S1C**), despite the comparable decrease in CO_2_-evoked freezing. This result is consistent with the lack of a robust jumping response to BNST acidosis and further supports a model in which the BLA, rather than the BNST, regulates CO_2_-evoked jumping responses.

Given the above results, we chose a different, more specific targeting strategy for disrupting ASIC1A in the amygdala. We hypothesized that BLA principal neurons may be the key site of ASIC1A action given that they are the most abundant neuron type in the BLA [32] and that these neurons have been implicated in fear- and anxiety-related behaviors [33]. Previous studies have used the CaMKII promoter to virally target these neurons [34–38]. Thus, here we injected an AAV expressing Cre recombinase under control of the CaMKII promoter (AAV-CaMKII-Cre) in *Asic1a^loxP/loxP^* mice to disrupt ASIC1A in BLA principal neurons. We found that this manipulation markedly decreased the expression of ASIC1A protein (**Figures 5A**, **5B**) suggesting that these neurons normally express ASIC1A protein. Moreover, targeting ASIC1A disruption with AAV-CaMKII-Cre reduced ASIC1A protein levels to an extent similar to that observed with AAV-CMV-Cre transduction (**Figures 4A**, **4B**). However, despite a similar reduction in ASIC1A protein, AAV-CaMKII-Cre had no effect on either jumping or freezing, in contrast to the reduction in freezing seen with AAV-CMV-Cre. Surprisingly, when exposed to 10% CO_2_, mice transduced with AAV-CaMKII-Cre behaved similarly to AAV-CaMKII-GFP-transduced controls and exhibited minimal jumping and a normal level of freezing (**Figure 5C**). Locomotion in the open field was similarly unaffected by AAV-CaMKII-Cre injection (**Figure S2B**). Together, these data suggest that in the BLA, ASIC1A may function in a different cell population, such as inhibitory interneurons or non-neuronal cells, to regulate CO_2_-evoked jumping. Notably, AAV-CaMKII-Cre also failed to reduce freezing, in contrast to the effect seen with AAV-CMV-Cre. This discrepancy could reflect a role for ASIC1A in non-principal neuron populations or non-neuronal cells in regulating freezing as well, though differences in viral transduction efficiency between the two constructs cannot be excluded.

**Figure 5.**
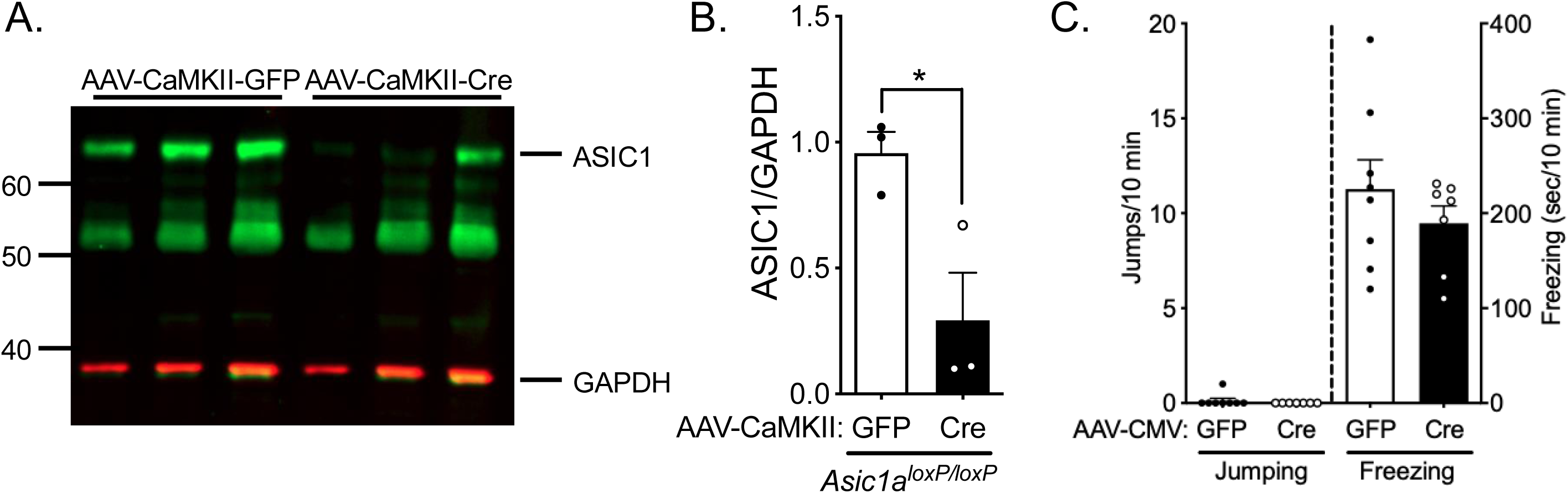
Disrupting ASIC1A in BLA principal neurons does not alter CO_2_-evoked defensive responses. ASIC1A was disrupted in BLA principal neurons of *Asic1a^loxP/loxP^* mice using an AAV expressing Cre recombinase (AAV-CaMKII-Cre) under control of a CaMKII promoter. An eGFP vector (AAV-CaMKII-GFP) was used as a control. **A, B)** AAV-CaMKII-Cre injection into the amygdala reduced ASIC1 protein (green) in amygdala punches as measured by western blotting. GAPDH (red) was used as a loading control. AAV-CaMKII-Cre transduction in the BLA reduced ASIC1A expression relative to AAV-CaMKII-GFP injected controls (t(4) = 10.24, *p = 0.0324, n = 3/group) **C)** Principal neuron-specific disruption of ASIC1A did not alter jumping (p > 0.9999, n = 8, 7) or freezing (t(13) = 0.9709, p = 0.3493) to 10% CO_2_.

We previously tested the effects of neuron-specific ASIC1A disruption on CO_2_-evoked freezing and found that mice with neuron-specific ASIC1A disruption (*Syn Asic1a* KO mice) displayed a reduction in CO_2_-evoked freezing similar to that observed in *Asic1a^-/-^*mice [24]. We therefore re-analyzed videos of Syn Asic1a KO mice to determine whether they also displayed CO_2_-evoked jumping. We observed increased CO_2_-evoked jumping in *Syn Asic1a* KO mice (**Figure S3**), supporting a neuronal site of action. This jumping was less pronounced than that observed with global ASIC1A disruption, which may reflect that in these mice some neuronally expressed ASIC1A protein remained intact [24, 28]. Our previous characterization found that Syn-Cre-mediated disruption was more complete in interneurons than in glutamatergic neurons, at least in cortex [24, 28]. Taken together with the lack of effect of CaMKII-Cre-mediated ASIC1A disruption, these observations raise the possibility that interneurons might be a key site.

To more directly assess how ASIC1A shapes neuronal responses to CO_2_ inhalation, we implanted microwire arrays into the BLA (**Figure S4**) and measured individual neuron firing and local field potentials (LFPs) during exposure to compressed air, 10%, and 20% CO_2_. We identified 83 single-unit waveforms in wild-type mice and 119 in *Asic1a^-/-^* mice. 10% CO_2_ predominantly suppressed firing, with only a minority of neurons showing increased firing (**Figure 6A)**. The distribution of responses was significantly different between genotypes in 10% CO_2_ (**Figure 6A**), with a higher proportion of *Asic1a^-/-^* neurons showing both increased and decreased firing, and fewer showing stable, unchanged firing. In 20% CO_2_, the proportions of suppressed neurons were greater and the proportions of excited neurons were smaller than in 10% CO_2_ in both genotypes. Though in 20% CO_2_ the distribution of responses did not differ significantly between genotypes, the distribution of responses was similar to that seen in 10% CO_2_.

**Figure 6.**
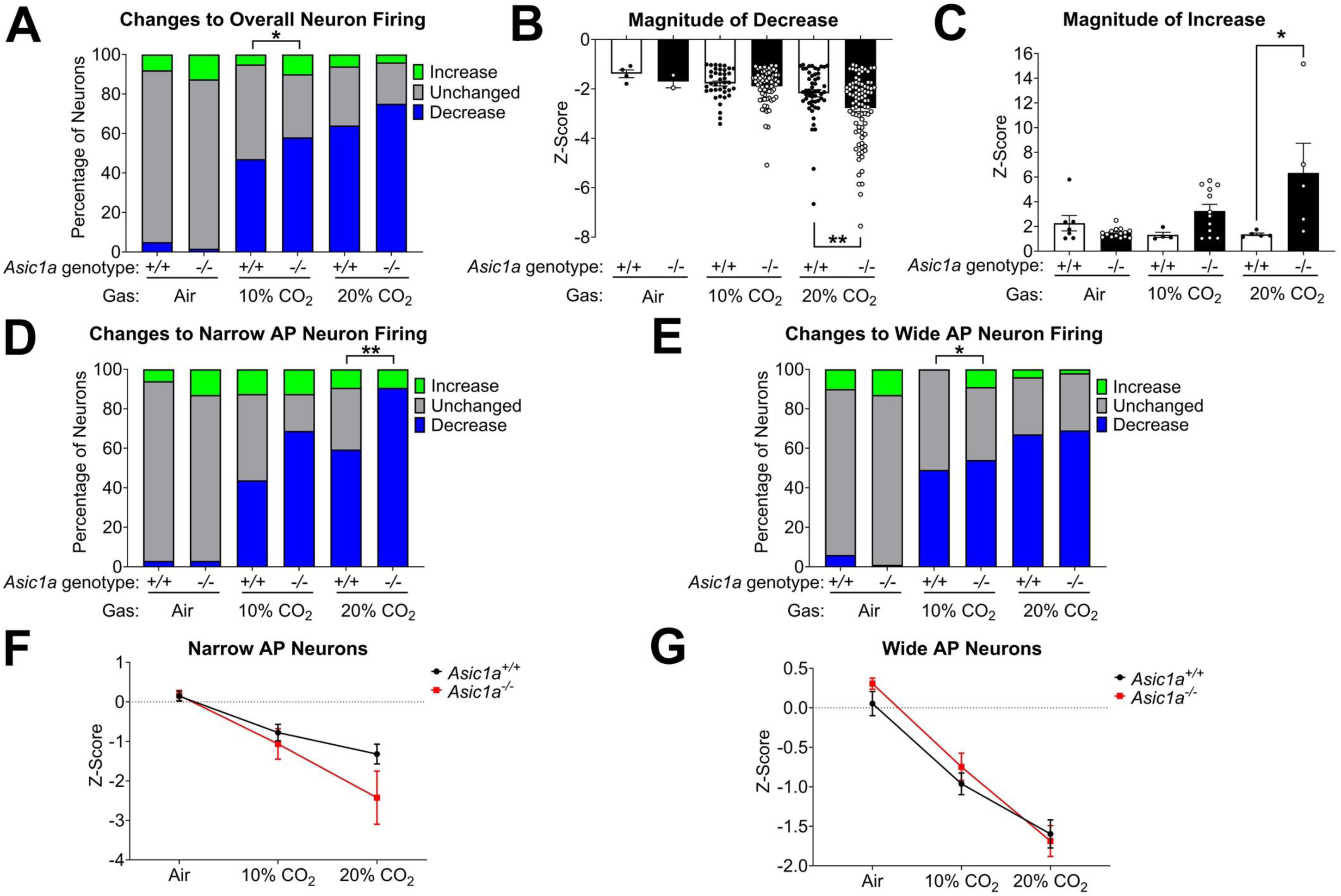
CO_2_ differentially affects BLA neuron firing. **A)** Distribution of neuronal responses to gas exposures. Firing frequency during gas exposure was z-normalized to baseline activity. Neurons with activity z < -1 were considered to have decreased firing, neurons with activity z > 1 were considered to have increased firing, and z-scores -1 < z < 1 were considered unchanged. Distribution of neuronal responses was significantly different between genotypes in 10% CO_2_ (χ*^2^* (df 2) = 6.165, p = 0.0459) but not in air (p = 0.263) or 20% CO_2_ (p = 0.247). **B)** Magnitude of activity change in neurons that decreased firing during gas exposure. In 20% CO_2_, *Asic1a^-/-^*neurons showed a significantly larger decrease in firing (U = 1729, **p = 0.008). In air and 10% CO_2_, wild-type and *Asic1a^-/-^* neurons showed similar magnitudes of decrease (Air: U = 2, p = 0.533; 10% CO_2_: U = 1222, p = 0.433). **C)** Magnitude of activity change in neurons that increased firing during gas exposure. In 20% CO_2_, *Asic1a^-/-^* neurons showed a higher magnitude of increased firing (U = 1, **p = 0.016). In 10% CO_2_, *Asic1a^-/-^* neurons showed a tendency toward higher magnitude of increased firing that did not reach significance (U = 10, p = 0.103). In air, wild-type and *Asic1a^-/-^* neurons showed similar magnitudes of increase (U = 41, p = 0.448). **D)** Distribution of neuronal responses only in neurons with narrow action potentials (putative interneurons). In this category, *Asic1a^-/-^* neurons differed from *Asic1a^+/+^* neurons in response to 20% CO_2_, with the majority of *Asic1a^-/-^*neurons decreasing firing and none staying unchanged (Fisher’s Exact Test p < 0.001). Distributions of responses to air and 10% CO_2_ did not differ between genotypes (Fisher’s Exact Test, Air p = 0.833; 10% CO_2_ p = 0.098). **E)** Distribution of neuronal responses only in neurons with wide action potentials (putative principal neurons). In this category, *Asic1a^-/-^*neurons differed from *Asic1a^+/+^* neurons in response to 10% CO_2_, notably showing some *Asic1a^-/-^*neurons increased firing whereas *Asic1a^+/+^*neurons did not (χ*^2^* (df 2) = 6.386, p = 0.041). Distributions of responses to air and 20% CO_2_ did not differ between genotypes (Fisher’s Exact Test, Air p = 0.296; 20% CO_2_ p = 0.835). **F)** Average response of individual neurons with narrow action potentials (putative interneurons) across gas conditions. There was a significant gas by genotype interaction (p = 0.045). **G)** Average response of individual neurons with wide action potentials (putative principal neurons) across gas conditions. No significant gas by genotype interaction was found in this subgroup (p = 0.68).

In addition to categorizing individual neurons’ responses to CO_2_ inhalation, we also measured the magnitude of each neuron’s change in firing. Among neurons that decreased their firing, the magnitude of suppression was greater in *Asic1a^-/-^* than in wild-type neurons in 20% CO_2_ (**Figure 6B**). Among neurons that increased their firing, the magnitude of excitation was greater in *Asic1a^-/-^* neurons in 20% CO_2_ (**Figure 6C**). This pattern was also observed in 10% CO_2_, though it did not reach significance.

Because our previous results suggested that interneurons rather than principal neurons might be the site of ASIC1A action, we further divided neurons into subtypes based on the width of their action potentials. The distribution of subtypes did not differ between genotypes (**Figure S5A**). In neurons with narrow waveforms (putative interneurons), ASIC1A disruption increased the proportion with suppressed firing (significant in 20% CO_2_, trending in 10% CO_2_; **Figure 6D**). In contrast, in neurons with wide waveforms (putative principal neurons), ASIC1A disruption increased the proportion with increased firing in 10% CO_2_ (**Figure 6E**). The decrease in putative interneuron firing and increase in putative principal neuron firing together contributed to the overall genotype difference in CO_2_ response distribution (**Figure 6A**). These results suggest that ASIC1A acts to maintain normal levels of interneuron firing during CO_2_ exposure, and that loss of this function may facilitate excitation of principal neurons.

The overall patterns seen for magnitude of neuron firing rate change were also seen when neurons were divided by action potential width (**Figure S5B-E**), and an overall analysis of neuron firing change across gases showed a statistically significant gas by genotype interaction in putative interneurons (**Figure 6F**) but not putative principal neurons (**Figure 6G**). These results further support the idea that ASIC1A plays a stabilizing role in BLA during CO_2_-induced acidosis, and that loss of ASIC1A results in disinhibition of the BLA, consistent with our earlier c-Fos results (**Figure 2**).

We further examined LFPs to determine how these neuron-level activity changes affected summated neural activity in the BLA. Whereas LFPs recorded in compressed air were similar to a baseline period during which no gas was infused into the chamber, CO_2_ inhalation rapidly and markedly altered LFP power (**Figure 7A**). In 10% CO_2_, LFP power was suppressed at most frequencies, though activation was seen in the theta (∼6 Hz) and high gamma (∼100 Hz) frequency ranges, which have been previously associated with fear memory [39–44]. 20% CO_2_ inhalation induced even greater suppression of LFP power, though activation in the theta range was still present. Interestingly, ASIC1A disruption resulted in stronger theta activation, enhanced high gamma activation, and greater suppression across low and mid gamma frequencies during inhalation of both 10% and 20% CO_2_ (**Figure 7B**). Together, these observations indicate that CO_2_ inhalation has profound effects on the BLA at both the neuron and circuit levels and that these changes depend in part on ASIC1A.

**Figure 7.**
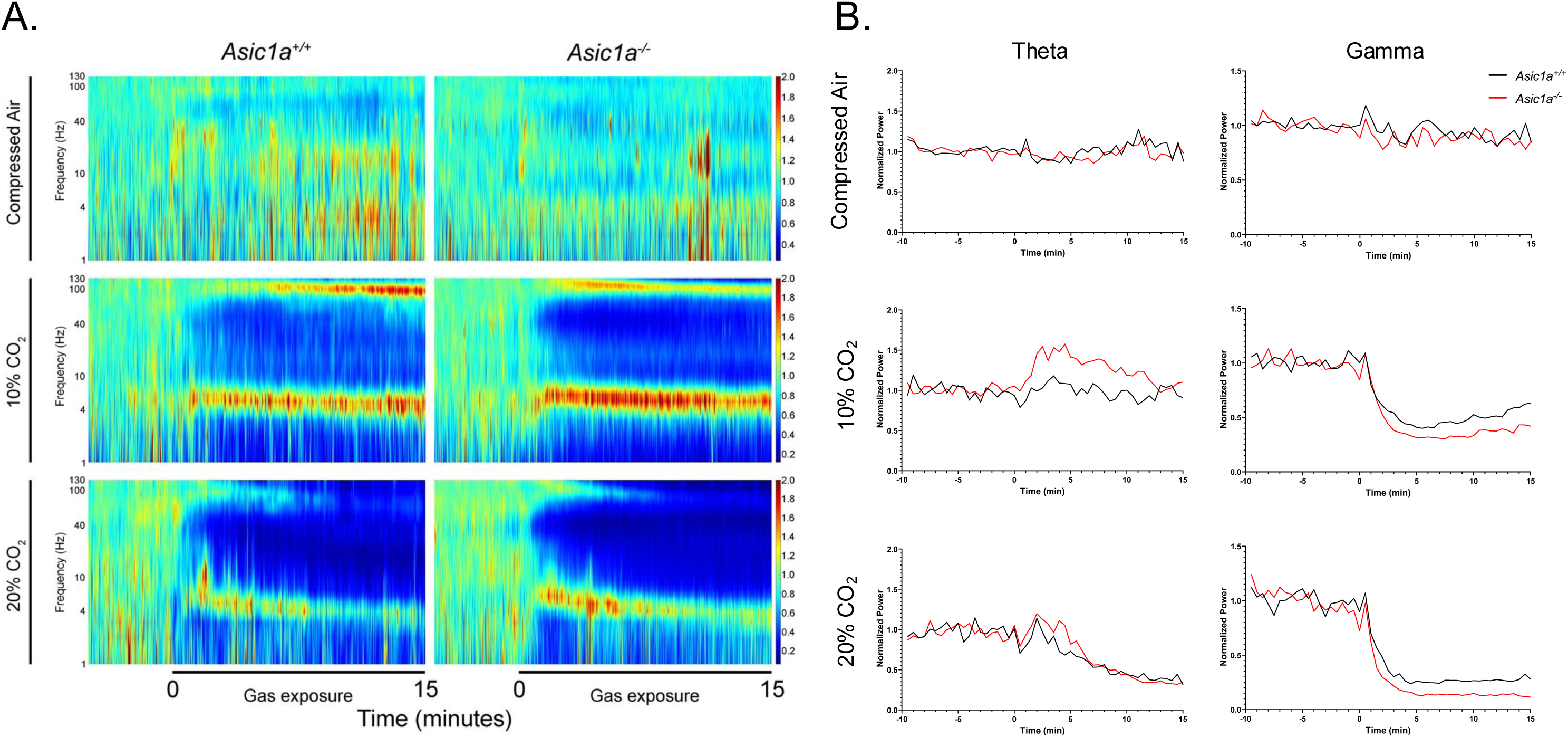
CO_2_ has prominent effects on BLA activity. **A)** Power spectrograms for *Asic1a^+/+^*and *Asic1a^-/-^* mice across exposures to various CO_2_ concentrations (compressed air, 10% CO_2_, and 20% CO_2_). Power is normalized to each respective pre-exposure baseline period (-8 to -2 minutes pre-exposure). 10% CO_2_ induces activation in the theta and high gamma frequency ranges (particularly near 6 Hz and 100 Hz) but causes suppression of power at other frequencies. In 20% CO_2_, power suppression is stronger, activation at 100 Hz is lost, and activation in the theta range is reduced. n = 7-9. Mixed-effects modeling of data from minutes 5-14 of compressed air and 10% CO_2_ exposure showed time*genotype*gas effects in delta (p = 0.0131), theta (p < 0.0001), alpha (p = 0.0001), beta (p = 0.0003), gamma (p = 0.0033), and high gamma (p < 0.0001). Genotype*gas effects were also seen in delta (p = 0.0030), theta (p < 0.0001), alpha (p < 0.0001), beta (p < 0.0001), and high gamma (p < 0.0001). **B)** Time course of field potential power over time for frequency bands of interest: theta (5-8 Hz) and gamma (32-48 Hz). *Asic1a^-/-^*mice showed stronger activation of theta and suppression of gamma to CO_2_ than *Asic1a^+/+^*mice.

## Discussion

Collectively, these findings support a key role for the BLA and ASICs in regulating defensive responses evoked by CO_2_ inhalation and brain acidosis [9–11, 13, 14, 24]. These data further demonstrate that ASICs regulate CO_2_-evoked jumping behavior as well as freezing, and that the two behaviors are dissociable. Moreover, the results shed important new light on the mechanisms underlying ASICs’ effects on these defensive behaviors. The observation that injecting acidic solution directly into the amygdala, but not the BNST, recapitulates both CO_2_-evoked freezing and jumping implicates CO_2_-induced acidosis as causal in driving these behaviors. This result is consistent with the known pH-sensitivity of ASIC1A and suggests that the amygdala is responding at least in part to local changes in pH.

An important finding of this study is that freezing and jumping are not simply inversely related opposing behaviors, but rather they are distinct and separable. Site-specific disruption of ASIC1A in the amygdala using CMV-Cre reduced CO_2_-evoked freezing without increasing jumping, with similar selective effects seen in BNST-specific disruption of ASIC1A. These observations indicate that the circuits controlling freezing and jumping are at least partially dissociable and suggest that the emergence of jumping in *Asic1a^-/-^* animals is not merely a consequence of reduced freezing. This dissociability may be functionally adaptive; jumping is likely to be more metabolically costly. Furthermore, jumping is more likely to draw attention (e.g. from predators) and thus would benefit from more stringent gating than freezing, which is comparatively lower cost and lower risk.

The dissociability of freezing and jumping is also supported by our data with ASIC2 disruption. We found that ASIC2 also contributes to CO_2_-evoked defensive responses, with ASIC1A and ASIC2 disruption producing comparable reductions in freezing, consistent with previous findings [14]. However, there were differences in jumping: both *Asic1a^-/-^* and *Asic2^-/-^*mice showed elevated jumping in 10% CO_2_, but only *Asic1a^-/-^* mice showed elevated jumping in 20% CO_2_. This difference may reflect differential expression of ASIC subunits across cell types and brain regions [14, 45], differences in channel physiology conferred by the two subunits [46, 47], or distinct neurophysiological consequences of disrupting each subunit [48]. The heightened jumping in both *Asic1a^-/-^* and *Asic2^-/-^* mice indicates that ASICs’ normal function is to oppose jumping, further implying that some other pH-sensitive molecule may serve as the acid sensor that drives jumping. Candidates include TRP channels, TASK channels, P2X receptors, and pH-sensitive G-protein-coupled receptors such as TDAG8, which has previously been implicated in CO_2_-evoked freezing [12, 49].

Our *in vivo* electrophysiological recordings represent, to our knowledge, the first characterization of how CO_2_ inhalation affects neural activity in the BLA in real time. Analysis of individual neuron firing showed that CO_2_ predominantly suppressed neuron firing in BLA, with some proportion of neurons unaffected, and a few increasing their firing. But the distribution of responses differed between genotypes: *Asic1a^-/-^* neurons showed a higher proportion of both increased and decreased firing in 10% CO_2_, and in 20% CO_2_, the magnitude of both increases and decreases was significantly greater in *Asic1a^-/-^* animals. Classifying neurons by action potential width, a proxy for cell type [30, 50], provided further information. Putative interneurons (narrow waveforms) in *Asic1a^-/-^* mice were more uniformly suppressed by 20% CO_2_ than those in wild-type mice, whereas putative principal neurons (wide waveforms) in *Asic1a^-/-^* mice were more likely to increase firing during 10% CO_2_. These findings suggest that ASIC1A normally acts to sustain inhibitory interneuron firing during CO_2_-induced acidosis, thereby maintaining tonic suppression of principal neuron output. When ASIC1A is absent, interneurons are more readily silenced by acidosis, disinhibiting principal neurons and shifting the balance of amygdala output toward excitation. This interneuron-centric model is further supported by our other results. First, c-Fos expression was elevated in the BLA of *Asic1a^-/-^*mice both at baseline and after CO_2_ exposure, consistent with reduced inhibitory tone and increased principal neuron activation, as principal neurons comprise the majority of neurons in the BLA [51]. The elevated c-Fos and jumping observed in *Asic1a^-/-^* mice in compressed air may also be consistent with this model, as the baseline condition is not affectively neutral — exposure to a novel environment and compressed air flow are themselves likely threatening albeit less than CO_2_, and loss of ASIC1A-dependent inhibitory tone may lower the threshold for BLA activation across diverse threats and intensities. Second, disrupting ASIC1A in BLA principal neurons (CaMKII-Cre) did not alter either freezing or jumping despite markedly reducing ASIC1A protein, suggesting that although principal neurons express ASIC1A, the behaviorally relevant ASIC1A activity was not in the principal neurons themselves.

These neuron level changes were also reflected at the scale of LFP changes in the BLA. CO_2_ produced a rapid, dose-dependent, and dramatic alteration of LFP power. In 10% CO_2_, LFP power was suppressed across most frequency bands, with notable exceptions in the theta (∼6 Hz) and high gamma (∼100 Hz) ranges. In 20% CO_2_, the broadband suppression was even more profound: theta activation persisted but was reduced compared to 10% CO_2_, and the high gamma activation observed at 10% was lost. The broadband suppression of higher-frequency LFP power was reminiscent of the effects of acute alcohol intoxication on LFPs in BLA [52] and may reflect a generalized impairment of local neural processing analogous to the EEG slowing seen in encephalopathic states such as delirium. In contrast, activation of theta and high gamma power is noteworthy, as gamma power has been shown to couple to theta [53] and both theta and gamma oscillations in the BLA have been repeatedly associated with fear learning, threat detection, and communication with upstream and downstream structures including the prefrontal cortex [39–44]. The activation of high gamma and theta power during CO_2_ may therefore reflect the prioritization of threat-relevant signaling even as other cognitive functions are impaired.

Critically, the LFP response to CO_2_ was ASIC1A-dependent. *Asic1a^-/-^* mice showed stronger theta activation and greater suppression of gamma power compared to wild-type mice during both 10% and 20% CO_2_ exposure. These genotype differences in field potential dynamics paralleled the behavioral differences between genotypes and suggested that the balance between theta activation and gamma suppression may be a neural correlate of the shift from freezing-dominant to jumping-dominant defensive responses. The reduced theta and high gamma activation when CO_2_ was increased from 10% to 20% in both genotypes may represent threat processing becoming overwhelmed by the stimulus and may also reflect the narrower behavioral difference seen in the two genotypes at this higher CO_2_ concentration. The convergence of the c-Fos, CaMKII-Cre, and *in vivo* electrophysiology data points to a population of BLA interneurons as a key site of ASIC1A action. The enhanced gamma suppression in *Asic1a^-/-^* mice is also consistent with this interpretation, as gamma-band activity is thought to depend heavily on local interneuron network activity [54, 55]. The enhanced gamma suppression may also explain the increased theta activation in *Asic1a^-/-^*mice, with diminished local inhibition resulting in unfiltered coordination with upstream sites via theta oscillations [56].

The circuit outside of the BLA that mediates CO_2_-evoked jumping remains to be defined. The BLA is known to receive inputs from multiple brain sites, with the prefrontal cortex (PFC) prominently implicated in fear learning [57]. In the case of CO_2_, interoceptive sites such as insula and thalamus may also contribute [58]. We speculate the role of the BLA may be to integrate signals from upstream sites to help select the most appropriate defensive response, consolidate it into memory, and communicate it to downstream effector sites. One leading candidate for an effector site downstream of BLA is the dorsal periaqueductal gray (dPAG). Electrical activation of dPAG evokes escape-like defensive responses in rodents [59–61], and our previous work suggested a role for the dPAG in CO_2_-evoked jumping [11]. Interestingly, the dPAG also has robust ASIC1A expression [62]. Other candidate structures include the parabigeminal nucleus [63], the intermediodorsal thalamic nucleus [64], and the paraventricular hypothalamus [65].

ASIC1A’s role in stabilizing neuronal activity in the BLA likely helps the BLA maintain its function in the face of threatening stimuli. Especially in the case of CO_2_ inhalation, ASIC1A’s acid-sensing function in the BLA may help weigh incoming signals about respiration and acidosis from upstream sites. Our model that ASIC1A stabilizes interneuron function in the BLA is consistent with an increasing appreciation that ASIC1A current density is particularly high in populations of interneurons, including in hippocampus and BLA [66, 67] and the suggestion that ASIC1A in interneurons contributes to seizure termination [68] and neurovascular coupling [28]. Loss of ASIC1A and resultant disinhibition of the BLA may result in impaired integration of upstream signals, shunting unfiltered information to downstream effectors, shifting responses toward more extreme defensive behaviors, which are more metabolically costly and behaviorally risky. This may be reminiscent of human subjects with amygdala lesions lacking anticipatory anxiety but showing exaggerated panic [13].

A limitation in interpreting these results is that although our site- and cell-type specific manipulations of ASIC1A were robust, they were not absolute. The behavioral effects of neuron-specific ASIC1A disruption (*Syn Asic1a* KO mice), which produced a milder jumping phenotype than global knockout, is consistent with either incomplete disruption in those animals [24, 28] or an additional contribution of non-neuronal ASIC1A. Similarly, residual ASIC1A protein after viral Cre-mediated disruption makes it impossible to fully rule out brain sites or cell types in which we did not see an effect. For example, AAV-CMV-Cre-mediated disruption of ASIC1A in BLA reduced freezing but did not increase jumping, raising the possibility that jumping may be more tightly gated than freezing and depend on a threshold level of ASIC1A disruption (perhaps in interneurons) that our manipulation did not achieve. Moreover, although the CaMKII promoter has been widely used to target BLA principal neurons, recent work suggests that CaMKII constructs may also have non-specific targeting [69] and, similar to CMV-Cre, may not have achieved a threshold level of ASIC1A disruption. Our electrophysiological classification of neuronal subtypes is also a limitation. Neurons were classified based on action potential width, a widely used method, although it does not provide definitive discrimination between neuronal subtypes. Also, in 20% CO_2_ too few neurons increased firing, thereby limiting statistical power to detect significant effects of ASIC1A. An important future direction will be to directly test the role of ASIC1A in BLA interneurons in CO_2_-evoked behavior.

Finally, these findings in mice may have translational relevance for human psychiatric conditions characterized by dysregulated defensive responses. Humans undergoing CO_2_ challenges report responses spanning the defensive spectrum from anxiety to fear to panic [6, 13, 70], and individuals with panic disorder show heightened CO_2_ sensitivity [4–8]. Although we cannot know how closely mouse jumping parallels human panic, both represent exaggerated defensive responses. Studies of single nucleotide polymorphisms in the human *ASIC1* gene have linked genetic variation in *ASIC1* to panic disorder, heightened CO_2_ reactivity, and increased amygdala volume and reactivity [71, 72]. Together with the present findings, these observations raise the possibility that targeting ASICs or brain pH could offer therapeutic benefit for disorders characterized by excessive defensive responses such as panic disorder and PTSD.

## Acknowledgements

ACC was supported by NIMH T32MH019113 and VA IK2BX006118. NSN was supported by the Juanita Bartlett Professorship. JAW was supported by NIMH R01MH113325, NIDA R01DA052953 and R01DA037216, VA IO1BX004440, and the Roy J. Carver Charitable Trust. The Zeiss LSM710 confocal microscope at University of Iowa Central Microscopy Research Facility was funded by the NIH (SIG grant, S10RR025439). We thank Chantal Allamargot for her recommendations and assistance. We thank Joel Geerling for allowing us to use his Olympus BX61VS microscope.

---

**Figure S1.**
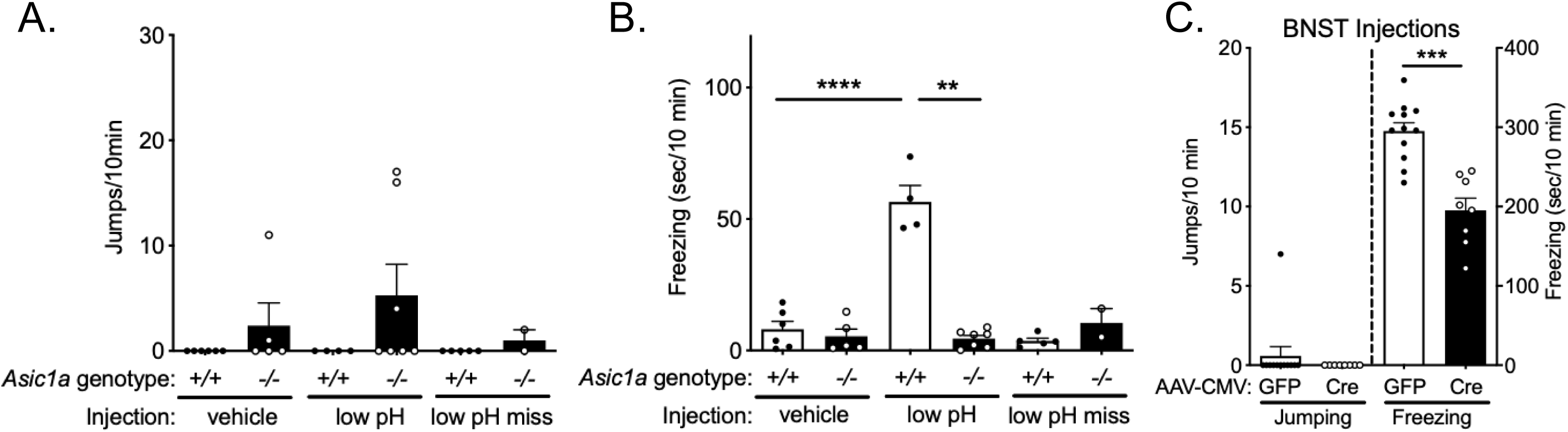
Acidification of the BNST elicits ASIC1A-dependent freezing and 10% CO_2_ Freezing depends on ASIC1A in BNST. **A)** Jumping behavior after infusion of low pH or normal pH solution into the BNST. Infusion of low pH into the BNST induced jumping behavior in some *Asic1a^-/-^* mice, though there was no treatment by genotype interaction (F(1,18) = 0.4303, p = 0.5201, n = 6, 5, 4, 7, 5, 2) and only a weak trend toward an effect of genotype (F(1,18) = 3.052, p = 0.0977). **B)** Freezing behavior after infusion of low pH or normal pH solution into the BNST. Infusion of low pH into the BNST induced ASIC1A-dependent freezing (*Taugher et al., 2014*). There was a significant treatment by genotype interaction (F(1,18) = 60.81, p < 0.0001). Wild-type mice injected with low pH solution displayed more freezing than *Asic1a^-/-^* mice injected with low pH (**p = 0.0029) or wild-type mice injected with a normal pH vehicle solution (****p < 0.0001). Seven low pH injections missed the BNST (5 wild-type, 2 *Asic1a^-/-^*); little freezing was observed in these mice. **C)** BNST-specific disruption of ASIC1A did not significantly alter jumping behavior (p > 0.9999, n = 12, 8), but did reduce freezing to 10% CO_2_ (Taugher et al., 2014) (t(18) = 5.532, ***p <0.0001).

**Figure S2.**
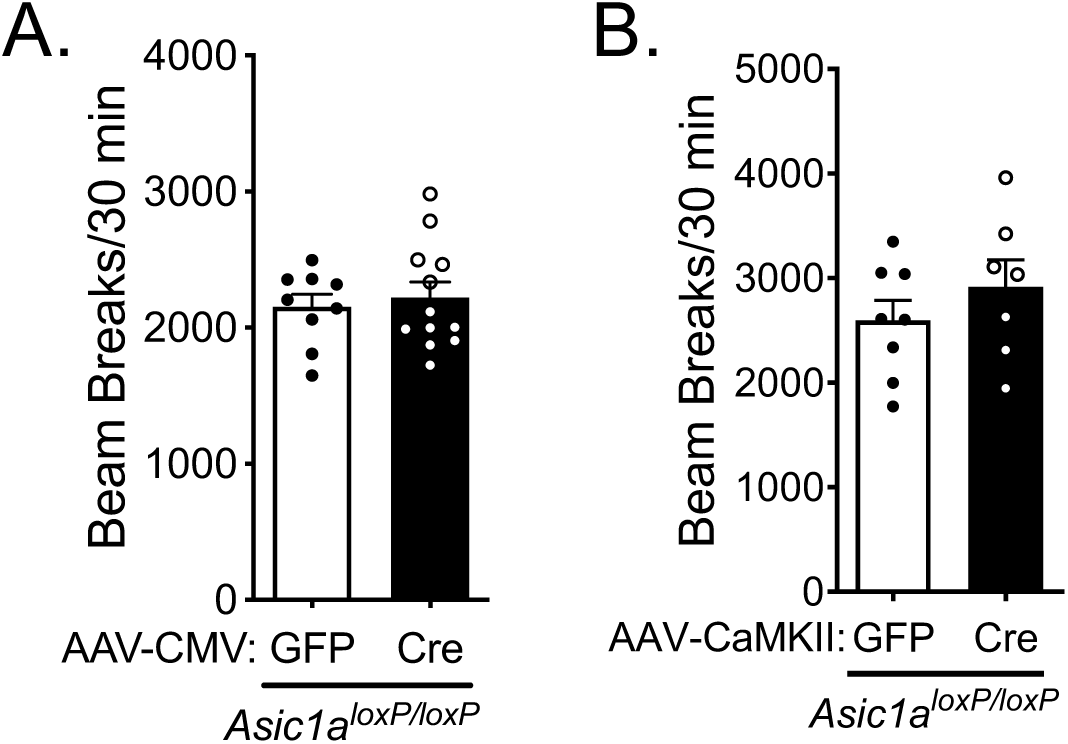
Disruption of ASIC1A in the amygdala with AAV-CMV-Cre or AAV-CaMKII-Cre does not alter locomotion in the open field. **A)** Locomotor activity in the open field was not affected by disruption of ASIC1A in the amygdala with AAV-CMV-Cre (t(19) = 0.4499, p = 0.6579). **B)** Similarly, ASIC1A disruption in BLA principal neurons with AAV-CaMKII-Cre did not alter locomotor activity in the open field (t(13) = 1.015, p = 0.3287).

**Figure S3.**
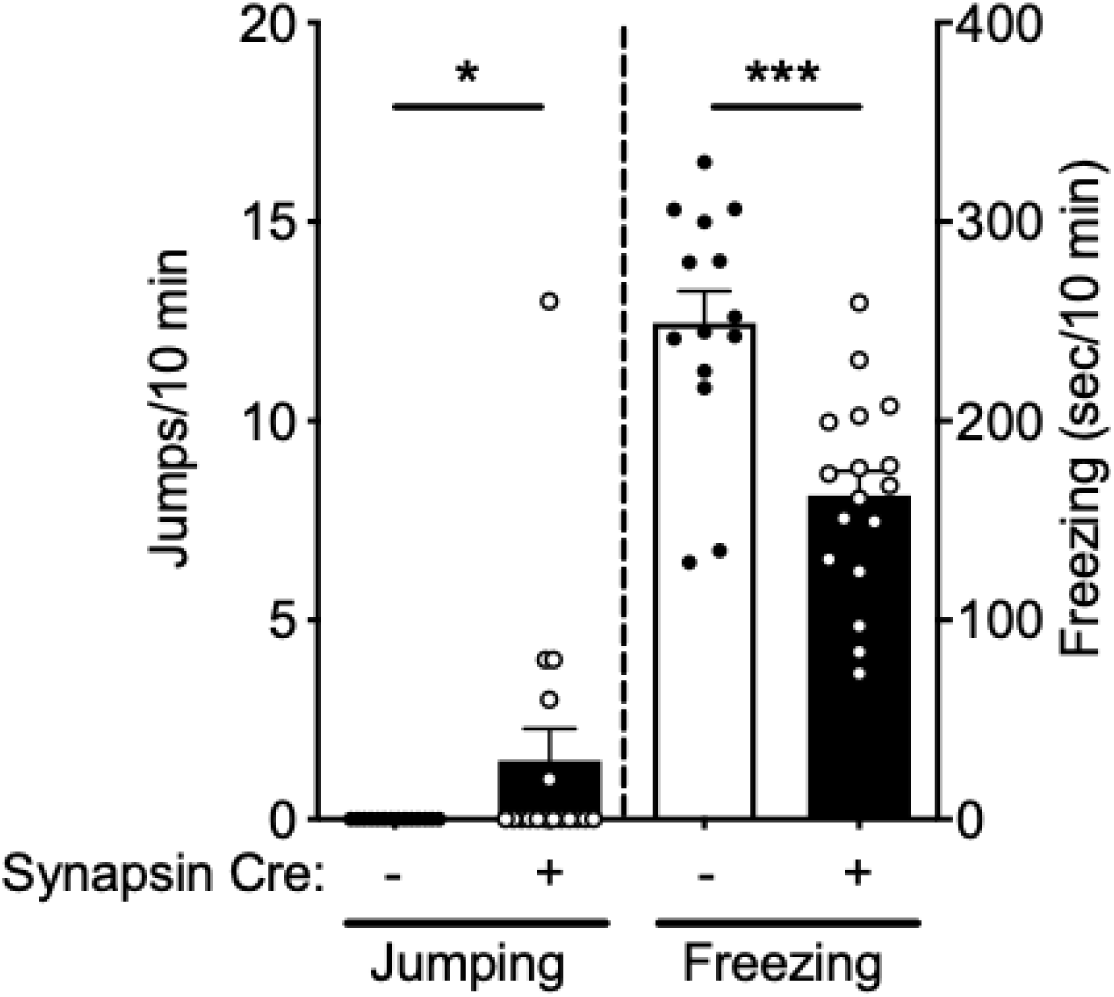
10% CO_2_ jumping and freezing depend on neuronal ASIC1A. Neuron-specific disruption of ASIC1A increased jumping in 10% CO_2_ (*p = 0.0482, n = 14, 17) and reduced freezing (Taugher et al., 2017) (t(29) = 4.365, ***p = 0.0001).

**Figure S4.**
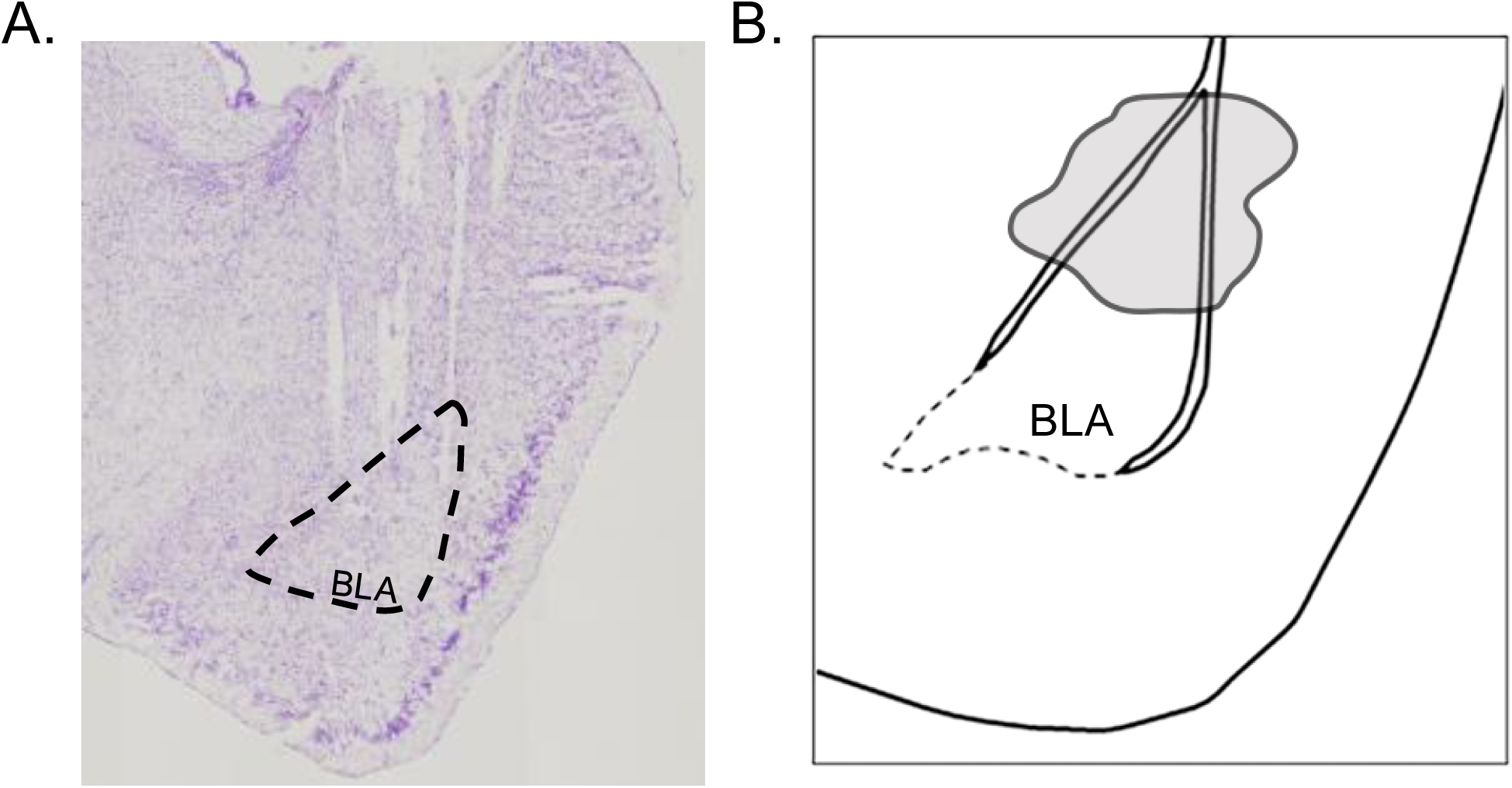
Microwire array placement. Placement of microwire arrays was verified postmortem. **A)** Representative Nissl-stained section showing tracts from wires within the BLA. **B)** Diagram of electrode placements. Implanted microwire arrays that hit the BLA were contained within the shaded area.

**Figure S5.**
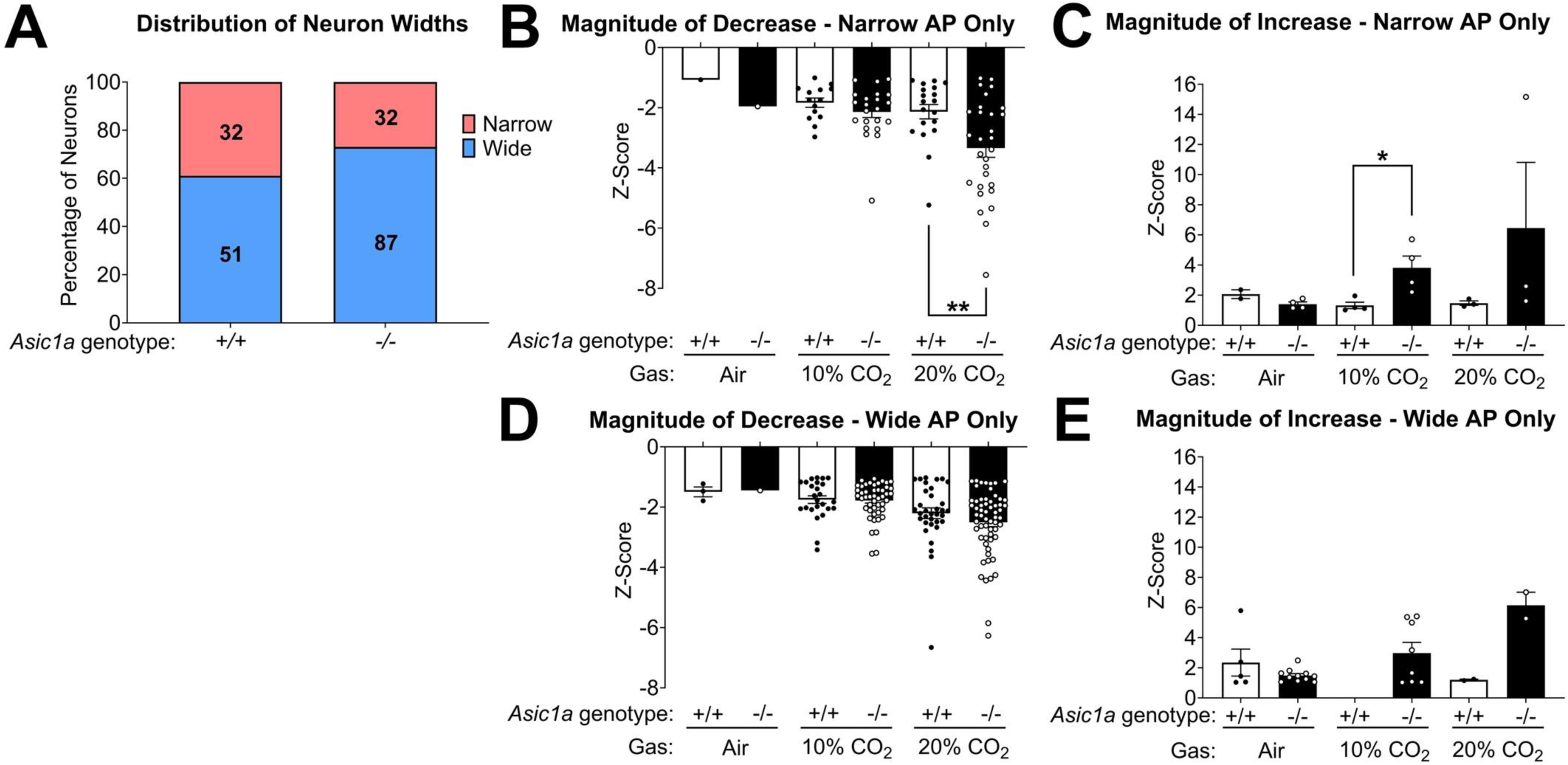
CO_2_ differentially affects BLA neuron types. **A)** Distribution of detected neurons with wide or narrow waveforms did not significantly differ between genotypes (p = 0.092). **B)** Magnitude of activity change only in neurons with narrow action potentials (putative interneurons) that decreased firing during gas exposure. In 20% CO_2_, *Asic1a^-/-^* neurons showed a significantly larger decrease in firing (U = 151, **p = 0.008). In 10% CO_2_, wild-type and *Asic1a^-/-^* neurons showed similar magnitudes of decrease (U = 119, p = 0.27). No testing could be done in air due to low n of neurons. **C)** Magnitude of activity change only in neurons with narrow action potentials that increased firing during gas exposure. In 10% CO_2_, *Asic1a^-/-^* neurons showed a higher magnitude of increased firing (U = 0, **p = 0.03). No statistically significant genotype difference was found in compressed air or 20% CO_2_ (Air: U = 0, p = 0.13, 20% CO_2_: U = 1, p = 0.20), but this subgroup analysis was underpowered. **D)** Magnitude of activity change only in neurons with wide action potentials (putative principal neurons) that decreased firing during gas exposure. No statistically significant genotype difference was found in 10% or 20% CO_2_ (10% CO_2_: U = 552, p = 0.68; 20% CO_2_: U = 854, p = 0.19). No statistical testing could be done in compressed air due to low n of neurons. **E)** Magnitude of activity change only in neurons with wide action potentials that increased firing during gas exposure. No statistically significant genotype difference was found in air or 20% CO_2_ (Air: U = 27, p = 0.1; 20% CO_2_: U = 0, p = 0.33). No testing could be done for 10% CO_2_ as no neurons fit this category in *Asic1a^+/+^* animals.

